# A linear mixed model approach to study multivariate gene-environment interactions

**DOI:** 10.1101/270611

**Authors:** Rachel Moore, Francesco Paolo Casale, Marc Jan Bonder, Danilo Horta, BIOS Consortium, Lude Franke, Inês Barroso, Oliver Stegle

## Abstract

Different environmental factors, including diet, physical activity, or external conditions can contribute to genotype-environment interactions (GxE). Although high-dimensional environmental data are increasingly available, and multiple environments have been implicated with GxE at the same loci, multi-environment tests for GxE are not established. Such joint analyses can increase power to detect GxE and improve the interpretation of these effects. Here, we propose the structured linear mixed model (StructLMM), a computationally efficient method to test for and characterize loci that interact with multiple environments. After validating our model using simulations, we apply StructLMM to body mass index in UK Biobank, where our method detects previously known and novel GxE signals. Finally, in an application to a large blood eQTL dataset, we demonstrate that StructLMM can be used to study interactions with hundreds of environmental variables.

## Introduction

Increasingly large population cohorts that combine genetic profiling with deep phenotyping and environmental information, including diet, physical activity and other lifestyle covariates, have fostered interest to study genotype-environment interactions (GxE). Already, such analyses have identified GxE for human phenotypes, including disease risk^1,2^, and molecular traits^3,4^.

Established methods to detect GxE implement tests that evaluate the effect of a single environmental variable on individual genetic variants^5^. Recent multivariate extensions enable assessing GxE across sets of genetic variants, either using genetic risk scores^6^ or variance component tests^7–9^.

Whilst there is evidence to suggest that multiple environments can interact with a single genetic locus to influence phenotypes, for example *FTO* interacts with a number of environments to alter BMI risk, including physical activity^10–13^, diet^12–15^ and smoking^12^, there exist no robust methods for the joint GxE analysis of multiple environmental variables. Such joint tests may increase the power to detect GxE, in particular when GxE effects are simultaneously driven by multiple environments, while at the same time reducing the multiple testing burden. Additionally, joint models of multiple environmental variables can improve the interpretation of GxE effects, allowing to assess the relevance of individual environments. As increasingly high-dimensional environmental data are available in population cohorts, there is a growing need for multi-environment GxE tests.

Here, we present a robust multi-environment GxE test based on linear mixed models (LMMs), using a random effect component to jointly model the effect of multiple environmental variables. The method generalises previous GxE tests to enable the joint analysis of hundreds of environmental variables and can be applied to large cohorts of hundreds of thousands of individuals.

## Results

Usually, LMMs test for persistent genetic associations of individual variants, that means constant genetic effect sizes across the population. Covariates and additional random effect components are included to account for population structure, environment, and other additive (confounding) factors. StructLMM extends the LMM framework by modelling heterogeneity in effect sizes due to GxE

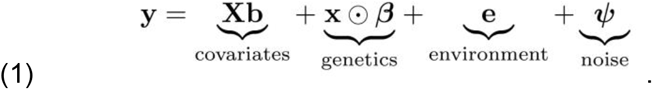

Here, *β* is a vector of per-individual effect sizes and follows a multivariate normal distribution

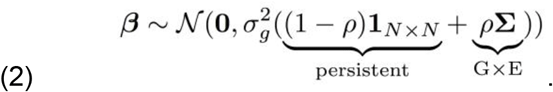

The first covariance term corresponds to a persistent genetic effect whereas the second covariance term accounts for heterogeneous effect sizes parameterised by an environmental covariance *Σ*, where the parameter ρ defines the fraction of genetic variance due to GxE. Depending on the functional form of *Σ*, this model can be used to account for different types of GxE, for example hierarchies of discrete environmental groups, or as considered here, GxE effects based on a set of continuous and discrete environmental covariates (**Fig. 1b-c**). The environmental covariance is also used to account for additive (i.e. non-genetic) environmental effects, *e* ~ *N*(0, *Σ*). The model is technically related to existing random effect tests for rare variants^16^ and epistasis^17^ (**Methods**).

**Figure 1.**
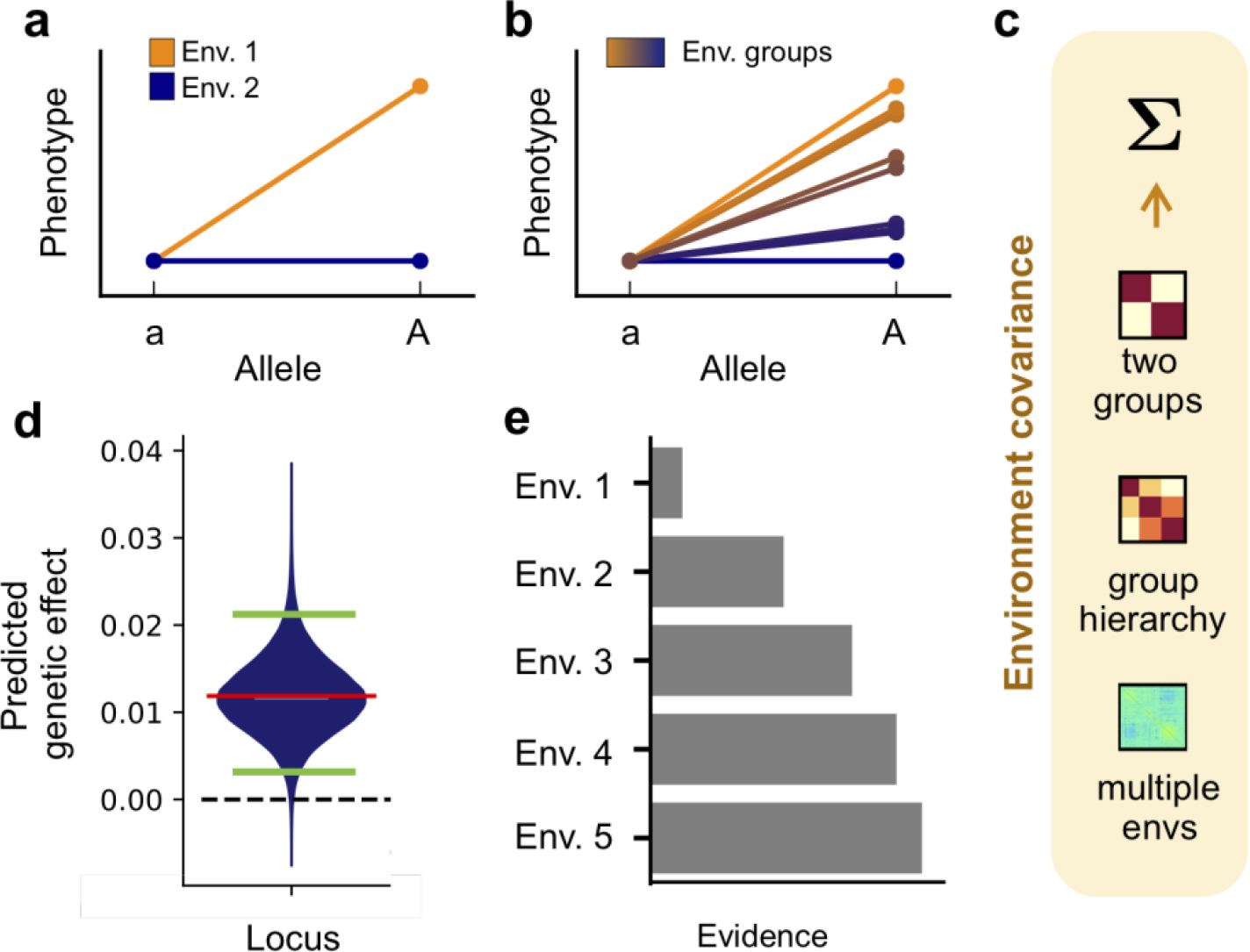
Overview of the StructLMM model. (**a**) Basic genotype-environment interaction, with a group-specific genetic effect (blue and orange lines correspond to the average phenotypes observed within two environmental groups for two alleles). (**b**) Interaction with multiple environmental groups or bins of continuous environmental states (average phenotypes for groups exerting increasing GxE effects from blue to orange for two alleles). (**c**) StructLMM accounts for heterogeneity in effect sizes due to GxE using a multivariate normal prior, where alternative choices of the environmental covariance *Σ* can capture discrete (two groups, group hierarchy; see **a**,**b**) or in the limit continuous substructure (multiple envs) of environmental exposures in the population. (**d,e**) Different example analyses using StructLMM. (**d**) Prediction of per-individual genetic effects in the population at individual loci. The violin plot displays the estimated density of individuals in a cohort that exert a genetic effect of a particular size given the distribution of the environmental factors within the population. Median and the top and bottom 5% quantiles of the effect size distribution are indicated by the red and green bars, respectively. (**e**) Evidence for the different environmental variables contributing to GxE effects.

Using the multi-environment model defined above (Eq. (1)), we propose score tests for two types of hypotheses: (i) an association test that accounts for heterogeneous effect sizes due to GxE and (ii) an interaction test to identify loci with significant GxE effects. StructLMM is computationally efficient, enabling genome-wide analyses using hundreds of environmental variables on cohorts of hundreds of thousands of individuals. The model facilitates different analyses to characterise GxE effects, including estimation of the fraction of genetic variance explained by GxE (ρ, Eq. (2)), and predicting per-individual genetic effect sizes based on environmental profiles in the population (**Fig. 1d**). Finally, the model can be used to assess the relevance of individual environmental variables for GxE (**Fig. 1e**). The full derivation of our method can be found in **Methods**.

## Model validation using simulated data

Initially, we considered simulated data using genotypes from the 1000 Genomes project^18^ to assess the statistical calibration and power of StructLMM. To mimic environmental distributions as observed in real settings, we simulated GxE based on 60 environmental covariates from UK Biobank, including physical activity, diet, and other lifestyle factors (**Methods**). We varied the sample size of the simulated population, the extent of GxE, the number of driving environments and other parameters (**Supp. Table 1**).

First, we confirmed the statistical calibration of StructLMM, considering phenotypes simulated without any genetic effects (i.e. the null model) (**Fig. 2a**, **Supp. Fig. 1a**) or simulated from a persistent effect model without interactions (**Supp. Fig. 1b**). Next, we simulated phenotypes with variable fractions of the genetic effects driven by GxE (ρ, Eq. (2)), and assessed power of the StructLMM association test (**Fig. 2b**). For comparison, we also considered a conventional LMM that ignores GxE, as well as a two-degrees-of-freedom (2-df) fixed effect model to jointly test for persistent associations and interactions using single environments (similar to the 2-df test in Kraft et al.^5^; Bonferroni adjusted for the overall number of environments, **Methods**). To facilitate direct comparisons, all considered models account for additive environmental effects using the same random effect term (Eq. (1)).

The power of the association tests decreased for larger GxE effects, demonstrating that strong GxE (large ρ) leads to reduced power for detecting associations (**Fig. 2b**). We also assessed power of StructLMM and single-environment fixed effect tests (one degree of freedom, e.g. Gauderman et al.^19^) for detecting GxE interactions based on the same data, where power increased for larger proportions of genetic variance due to GxE (**Fig. 2c**). Notably, both the StructLMM association and interaction tests were substantially better powered than existing methods, indicating that the model is broadly applicable to account for GxE. As a second parameter, we simulated phenotypes with increasing number of environments contributing to GxE effects, but tested for GxE effects using all 60 environmental variables. The results of this analysis show that StructLMM increasingly outperforms the corresponding single environment GxE model as the number of environments with non-zero GxE increases (**Fig. 2d, e**), in particular when testing for interaction effects (**Fig. 2e**).

**Figure 2.**
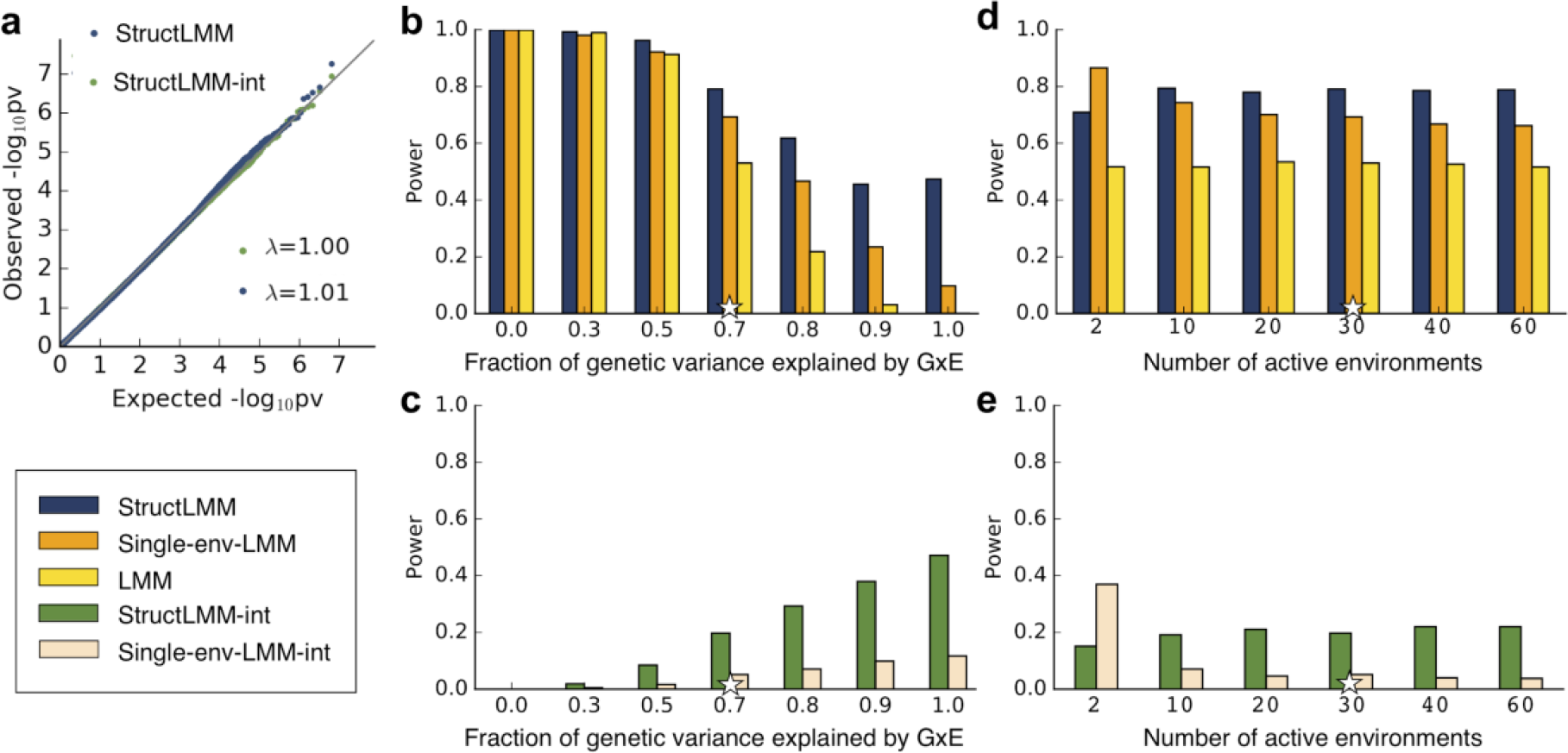
Assessment of statistical calibration and power using simulated data. (**a**) QQ plots of negative log P values from the StructLMM association test (blue, StructLMM) and interaction test (green, StructLMM-int) using synthetic data simulated from the null (no genetic effect). (**b**) Comparison of power for detecting genetic associations for increasing fractions of genetic variance explained by GxE (ρ). Included in this comparison were StructLMM, a 2-df fixed effect tests that jointly tests for persistent associations and interactions with a single environment (Single-env-LMM), as well as a conventional LMM to test for persistent effects (LMM). (**c**) Assessment of alternative methods for detecting simulated GxE effects using the same data and settings as in **b**. Compared were the StructLMM interaction test (StructLMM-int) and a single-environment interaction test (Single-env-LMM-int). (**d, e**) Analogous power analysis for detecting associations and interactions respectively, when simulating GxE using increasing numbers of environments with non-zero GxE effects (out of 60 environments total, considered in all tests). Models were assessed in terms of power (FWER<1%) for detecting simulated causal variants (**Methods**). Stars denote default values of genetic parameters, which were retained when varying other parameters (**Supp. Table 1)**.

We considered a number of additional settings, including varying the total number of observed environments, the fraction of phenotypic variance explained by additive environmental effects and additional forms of model mismatch, where phenotypes are simulated with interaction effects from environments that are not included at the testing stage. Across settings, StructLMM produced calibrated P values (**Supp. Fig. 2**) and had consistent power advantages over alternative methods (**Supp. Fig. 3**). Finally, we note that multi-environment GxE tests can in principle also be implemented based on fixed effect tests with as many degrees of freedoms as environments. However, we observed that such tests were not always calibrated and had lower power in some settings (**Supp. Fig. 4**), demonstrating the advantages of the random effect approach taken in StructLMM.

Taken together, these results show increased power and robustness of StructLMM compared to existing methods, in particular when large numbers of environments drive the GxE interaction effects.

## Application to data from UK Biobank

We tested for associations between low-frequency and common variants (imputed variants, MAF>1%, 7,515,856 variants in total) and BMI, considering 64 lifestyle covariates similar to those used in Young et al.^13^ (12 diet-related factors, three factors linked to physical activity and six lifestyle factors, modelled as gender-specific and age-adjusted, **Methods, Supp. Fig. 5, Supp. Fig. 6**) to account for GxE. A set of 252,188 unrelated individuals of European ancestry, for which all 64 environmental covariates, and the BMI phenotype were available in the full release of UK Biobank^20^ were taken forward for all analyses.

Initially, we applied a conventional LMM and StructLMM to test for associations, using an environmental covariance estimated using the 64 lifestyle factors to account for additive environmental effects in both methods and heterogeneity of genetic effects due to GxE in StructLMM. StructLMM and LMM identified slightly different sets of loci (323 and 327 loci were found by StructLMM and LMM, respectively), with 13 and 17 loci identified exclusively by StructLMM and LMM respectively (**Fig. 3a**, **Supp. Table 2**). As expected, the LMM had better detection power for loci with little or no GxE (**Supp. Fig. 12**). However, the distribution of ρ suggests substantial heterogeneity in genetic effect sizes for significant loci (**Supp. Fig. 12**), in particular for the 13 loci that were exclusively detected by StructLMM (**Supp. Table 2, Supp. Fig. 8-11)** (0.23<ρ<0.94). Additionally, StructLMM yielded lower P values for associations recovered by both models (**Fig. 3a**, e.g. P=1.57×10^−183^ vs P=1.84×10^−150^ for the well-documented *FTO* locus *rs1421085*), while retaining statistical calibration (**Supp. Fig. 7**).

Several of the additional loci identified by StructLMM have been previously associated to BMI or BMI-related traits. These included a variant (*rs11259931*) in *ADAMTSL3*, which codes for a glycoprotein^21^. The same variant and variants in LD (r^2^>0.53) have previously been linked to BMI-related traits, including lean body mass^22^, waist circumference^23^, hip circumference adjusted for BMI^24^ and height^25^. A second example is *rs11880064* in *PEPD*, which encodes for a protein involved in the final stage of degradation of endogenous and dietary proteins. Several additional *PEPD* genetic variants have been associated with adiponectin^26,27^, fasting insulin adjusted for BMI^28^, HDL cholesterol^29^, triglycerides^29,30^, type 2 diabetes^31^, waist circumference adjusted for body mass^24^ and waist to hip ratio^24^. A third association (rs473428) was identified upstream of *ONECUT1* and downstream of *WDR72*. *ONECUT1* stimulates the production of liver expressed genes and can inhibit glucocorticoid-stimulated gene transcription^32^ and genetic association with BMI^33^, cholesterol HDL^30^, lipids^30^ and triglycerides^30^ were reported in early GWAS results but have not reached genome-wide significance in more recent meta-analyses^34^, which may be due to GxE effects varying across the different aggregated cohorts or due to differences in trait transformation. Variants in *WDR72* are also associated to a large of number of relevant traits^30,33^.

**Figure 3.**
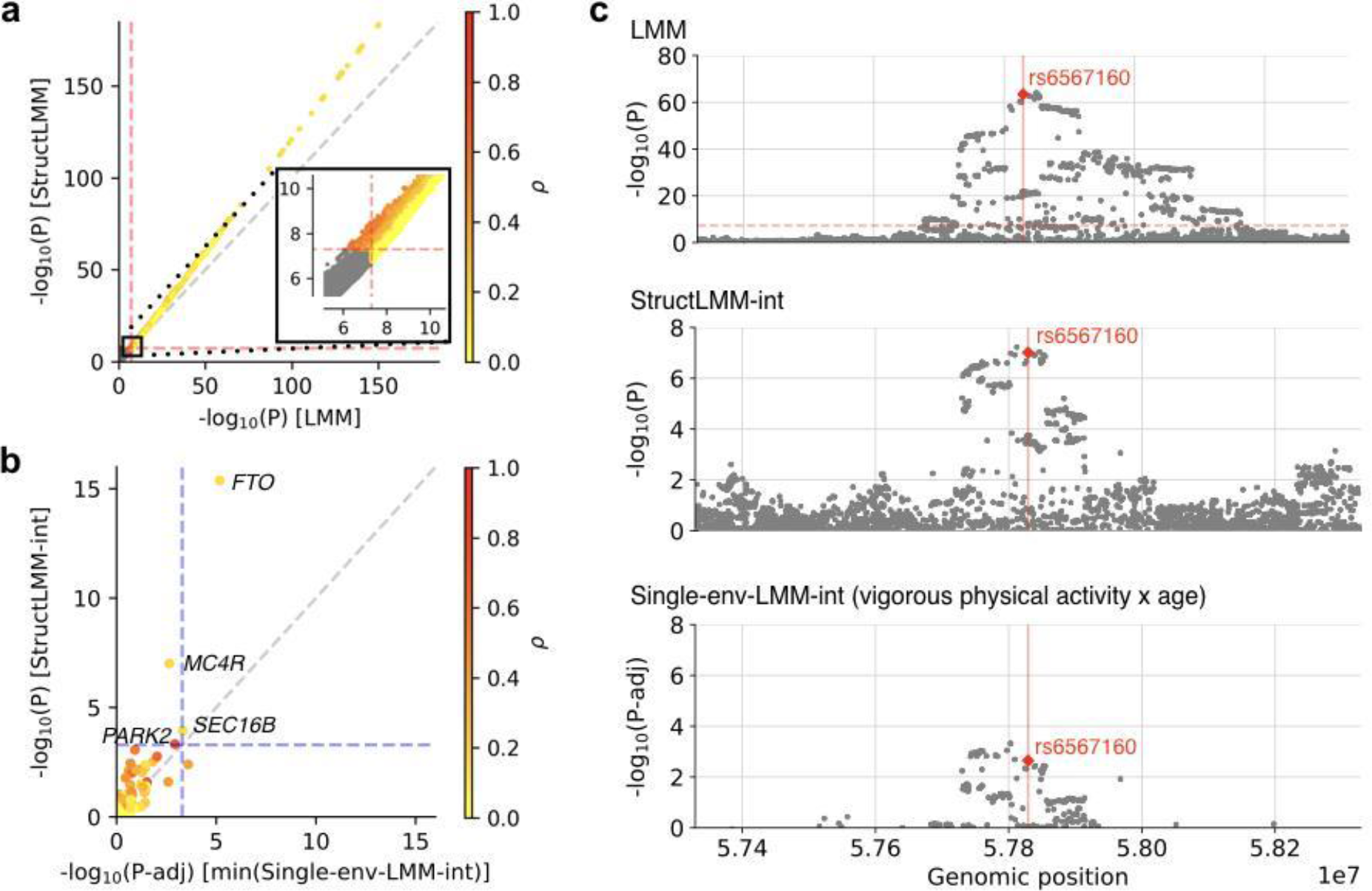
Applications to model GxE on body mass index (BMI) in UK Biobank. (**a**) Scatter plot of genome-wide negative log P values from a standard LMM (x-axis) versus the StructLMM association test (y-axis). Dashed lines indicate genome-wide significance at P<5×10^−8^ and colour denotes the estimated extent of heterogeneity (fitted parameter ρ), where yellow/red denotes variants with low/high GxE component. The inset displays a zoom-in view of variants close to genome-wide significance. (**b**) Scatter plot of negative log P values from GxE interaction tests at 97 GIANT variants^34^, considering a single-environment fixed effect GxE tests (x-axis, P values Bonferroni adjusted for the number of tested environments) versus the StructLMM interaction test (y-axis). Dashed lines correspond to alpha<0.05, Bonferroni adjusted for the number of tests. (**c**) Local Manhattan plots of an interaction identified by StructLMM at *MC4R*. From top to bottom: LMM association test, StructLMM interaction test, single-environment LMM interaction test for the environment with the strongest GxE effect at the GIANT SNP (vigorous physical activity x age). The red vertical line and diamond symbol indicates the GIANT SNP as in **b**.

Next, we applied StructLMM to test for GxE interactions. To reduce the number of tests, we tested for interactions at 97 GIANT variants (corresponding genes as annotated by GIANT^34^) that have previously been linked to BMI using independent data^34^. For comparison, we also applied a one degree of freedom fixed effect GxE test using individual environments (**Fig.,3b**, **Supp. Fig. 7, Supp. Fig. 14, Supp. Table 3**). Notably, StructLMM identified four significant GxE effects (α<0.05, Bonferroni adjusted), two of which were missed by the single-environment fixed effect test. Among the loci identified was the *FTO* locus (rs1241085, *ρ*ρ =0.14, **Supp. Fig. 14a**), which has previously been implicated with GxE for multiple environments^10,12–14^, *MC4R* (**Fig. 3c**) for which an interaction with physical activity in females aged 20-40yrs has been previously suggested (P-adj=0.025)^12^, *SEC16B* (**Supp. Fig. 14b**), for which secondary analyses provided some evidence for an interaction (P=0.025) with physical activity in Europeans^11^ and in a separate study in Hispanics^35^ and *PARK2* (**Supp. Fig. 14c**), a gene that has been linked to time-dependent variation in BMI suggested to be due to changes of environmental exposures^36^. StructLMM also enhanced the significance of test for interactions identified by both models, with significance levels for *FTO* and *SEC16B* dropping from P=4.23×10^−16^ to P-adj=6.76×10^−6^ and P=1.15×10^−4^ to P-adj=4.48×10^−4^ respectively, when considering a single-environment test. Larger differences in the number of discoveries were observed for more lenient threshold, e.g. 11 versus six loci with GxE at 5% FDR (Benjamini-Hochberg adjusted, **Supp. T6able 3**). Finally, we compared to a multi-environment GxE test based on fixed effects, which although calibrated on this large dataset, was underpowered (**Sup. Fig. 15**).

StructLMM can be used for the interpretation of GxE interactions, and in particular to predict per-individual genetic effects based on environmental profiles (**Fig. 4a**). We assessed the consistency of these estimates using hold-out validation, confirming that StructLMM can be used to explain and predict inter-individual variations in genetic effects due to GxE (**Supp. Fig. 16**). To identify environmental variables that drive the GxE signal, we used backward elimination, calculating Bayes factors between the full model and models with increasing numbers of environments removed. This analysis identified approximately 21 environmental factors that contribute to the GxE effect at *MC4R*, including all three physical activity measures for females, in agreement with^12^, but also identified a number of additional environments (some of which were more relevant) (**Fig. 4c**), underlining the benefits of multivariate modelling of GxE using sets of environments.

For other loci, we consistently observed that multiple environments contribute to GxE but there are large differences in the GxE architecture, with *FTO* driven by the largest number of environments (approximately 35) whilst *SEC16B* and *PARK2* were driven by a much smaller number of environments (approximately 9 and 10 respectively) (**Supp. Fig. 14d-f**). We also note that many of the environmental effects were gender specific for *MC4R* and *PARK2* and age dependent at *SEC16B*, with substantial overlap between the sets of interacting environments for three of the four loci, *PARK2* being the exception. Differences in the environments that drive these GxE effects were also apparent when correlating effect size predictions across loci, which identified groups of variants with similar GxE profiles (**Supp. Fig. 17**).

**Figure 4.**
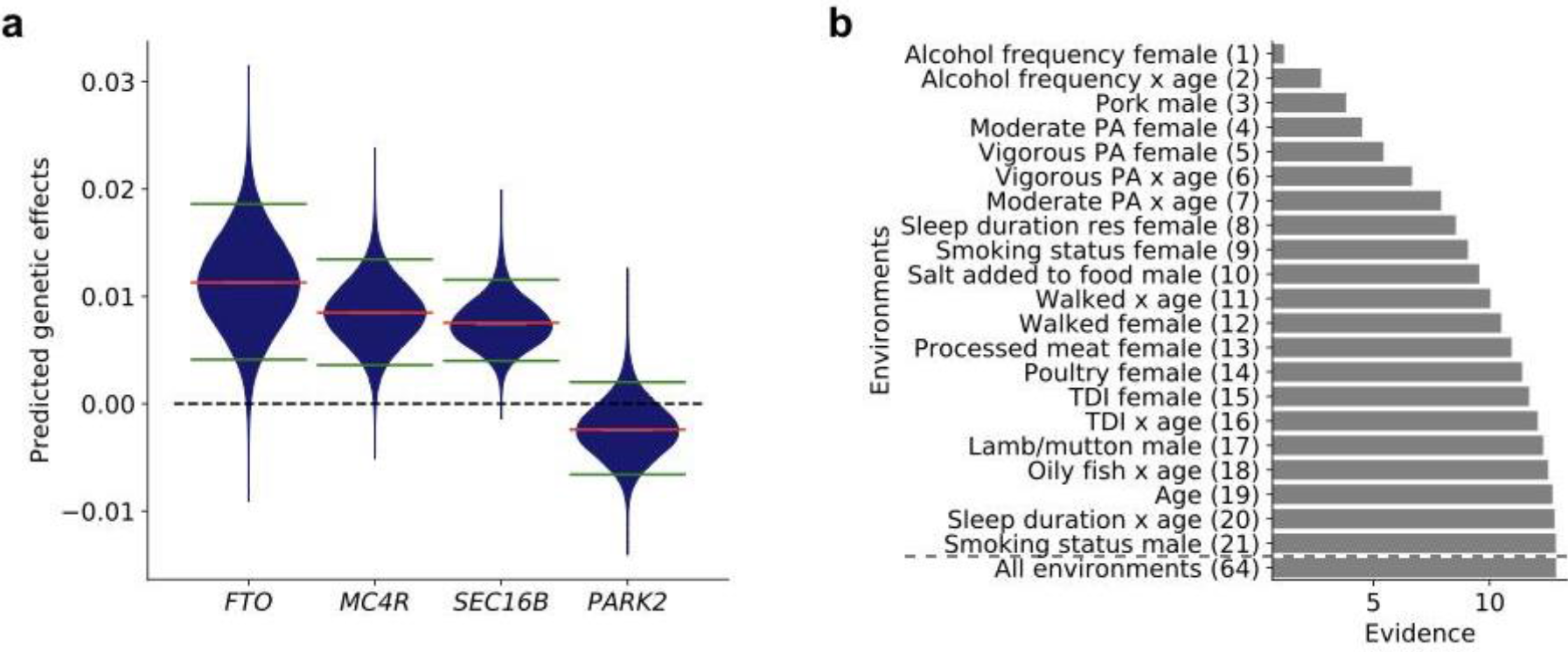
Analysis of environmental factors that drive GxE for BMI. (**a**) Violin plots showing distributions of estimated genetic effects on log BMI for the four GIANT variants with GxE (alpha<0.05, **Fig. 3b**). Estimated persistent genetic effects are shown by the red bar and the green bars indicate top and bottom 5% quantiles of variation in effect sizes due to GxE. (**b**) Cumulative evidence of environmental variables for GxE in order of relevance at *MC4R*, showing Bayes factors between the full model and models with increasing numbers of environmental variables removed using backward elimination. For comparison, shown is the total evidence of all environmental variables. ‘Alcohol frequency female’, is identified as the most important environmental factor, followed by ‘Alcohol frequency x age’ and so on.

## Identification of eQTL interactions with cellular state

As a second application, we considered a gene expression dataset and used StructLMM to identify regulatory variants with genetic effects that depend on cellular contexts, such as cell type^37^ or external stimuli^4^. The identification of such context-dependent genetic effects can help elucidate the regulatory mechanisms of disease loci by identifying relevant cell types and molecular pathways^38–40^.

We reanalyzed the large whole-blood expression dataset comprising of over 2,000 genotyped samples profiled with RNA-seq^41^ (**Methods**) and investigated cell-context interactions of *cis* expression quantitative trait loci (eQTL). Following^41^, we considered gene expression levels both as phenotypes but also as proxy variables to capture inter-sample variation due to changes in blood cell composition and other factors. Specifically, we considered a set of 443 highly variable genes as environmental factors in our analysis (**Methods**).

Initially, we applied a standard LMM to map *cis* expression quantitative trait loci (eQTL, plus or minus 250 kb from the centre of the gene, **Methods**). Next, we applied StructLMM to test for cell-context interactions at lead variants for 23,277 genes with an eQTL (FDR<5%, **Methods**). This identified 3,483 eQTL with a cell-context interaction (FDR<5%, **Supp. Table 4**), where StructLMM yielded calibrated P values despite the large number of environments (**Supp. Fig. 18**). Although overall, interactions with cell-context tended to explain only a small proportion of the *cis* genetic effect on gene expression variance (*ρ* < 0.2., for 68.0% of eQTL with a cell-context interaction, **Fig. 5a**), our analysis identified 532 genes (15.3% of eQTL with cell-context interaction) for which heterogeneity explained a larger proportion of the *cis* genetic variance than persistent effects (ρ> 0.5, **Fig. 5a**). As alternative method to detect cell-context interactions, we also considered multi-environment interaction tests based on fixed effects (**Methods**), which were less robust and identified fewer interaction eQTL than StructLMM (**Supp. Fig. 18**). Finally, we compared our interaction results to the findings of the primary analysis of the data^41^, where a step-wise procedure was employed to identify interaction eQTL (**Methods**). For 17,952 genes that were analysed in both studies, StructLMM identified 3,372 interaction eQTL compared to 1,841 interaction eQTL in the primary analysis (overlap 1,071, FDR<5%, **Supp. Fig. 18**).

**Figure 5.**
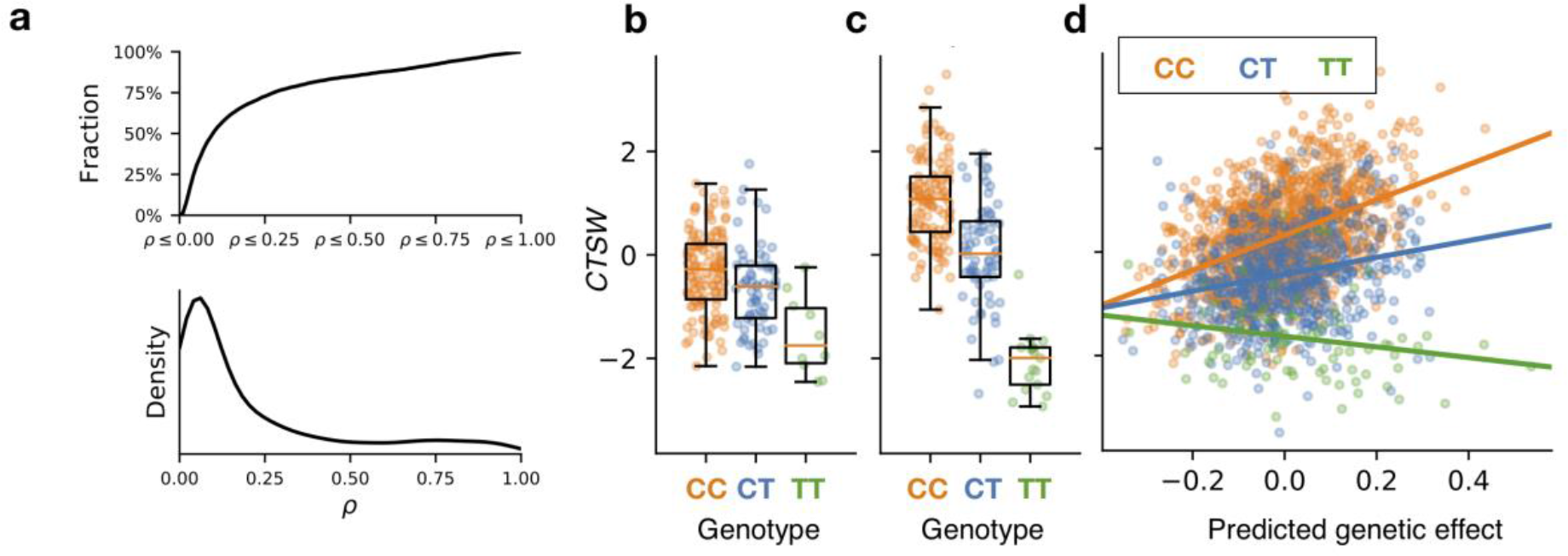
Analysis of gene-context interactions in a large blood gene expression cohort. (**a**) Cumulative fraction (top) and density (bottom) of eQTL with significant interactions as a function of the estimated degrees of heterogeneity (*ρ*). (**b**) Example of an interaction eQTL for *CTSW*, which is in LD with a risk variant for Crohn’s disease *rs568617* (r^2^=1, **Supp. Fig. 20**). (**b,c**) expression level of *CTSW* for different eQTL genotype groups in the 10% stratum of samples with the lowest (**b**) and highest (**c**) predicted genetic effect; analogous figure for the 10% strata. (**d**) *CTSW* expression level versus estimated per-individual genetic effects, stratified by the eQTL genotype. Analogous analyses for all 64 interaction eQTL with evidence for colocalisation with disease loci are provided in **Supp. Dataset 2**.

Next, we overlapped eQTL with cell-context interactions and risk variants from the NHGRI-EBI GWAS catalog V1.0.1^42^, which identified 64 instances of putative colocalisation (r^2^>0.8 of lead eQTL and GWAS variants, **Methods**), including GWAS variants for autoimmune diseases, infectious diseases and blood cell traits (**Supp. Table 5, Supp. Dataset 1)**. Notably, 46 of these eQTL with interactions were not reported in the primary analysis^41^. One of these is an interaction eQTL for *CTSW* expression (**Fig. 5c**, P=2.2×10^−15^, ρ=0.12), which is in LD with a risk variant for Crohn’s disease *rs568617* (r^2^=1.00, **Supp. Fig. 19**). To investigate the molecular pathways that drive this interaction eQTL, we stratified the population into groups with increased/decreased genetic effects predicted using StructLMM, and tested for enriched pathways among genes that were differentially expressed between these groups (**Methods**). This identified *positive T cell selection (GO:0046632)* and *positive regulation of interleukin-17 secretion* (*GO:0032740*) as cell-context environments that underlie the interaction eQTL (See **Supp. Table 5** for genome-wide enrichment results). Consistent with this, IL-17 producing CD4^+^ T cells are known to play a key role in the pathogenesis of inflammatory bowel disease, including Crohn’s disease43.

Taken together, the analysis of context-specific QTL using molecular traits demonstrates how StructLMM can be used to identify genetic effects on molecular traits that depend on the cellular context, where the model again outperformed existing methods. These results demonstrate the broad applicability of the model, including settings with large numbers of environmental factors.

## Discussion

We proposed a method based on LMMs to flexibly model GxE using sets of environments, thereby enabling the analysis of genotype-environment interactions with multiple environments. Conceptually, our approach is related to set tests for groups of variants, and offers power advantages when multiple environmental factors contribute to GxE (**Fig. 2**).

We applied StructLMM to data from UK Biobank, where the model detected association signals that were missed by an LMM, in particular when a substantial fraction of the genetic variance was explained by GxE. This demonstrates that accounting for heterogeneity in effect sizes (GxE) is not only of interest for mechanistic characterization of known genetic effects across environments, but can in some instances also increase the power to detect new genetic effects, which is similar to previous uses of 2-df fixed effect tests^5^.

When assessing GxE at 97 GIANT variants associated with BMI, we confirm established GxE effects for *FTO*, and we identified, for the first time, three additional GxE signals at stringent thresholds, some of which confirm prior evidence^11,12,14,15,35,36^. FDR-based significance, as frequently employed for GxE analyses^6,12^, would increase the number of discoveries further, yielding up to 11 GIANT variants with evidence for GxE on BMI (**Supp. Table 2**). In addition to offering power advantages, StructLMM yields per-individual predictions of variation in genetic effects due to GxE. We have shown that this allows for important downstream analyses, including the identification of individuals with increased or decreased genetic effects at different loci based on their own environmental exposure, and the identification of environmental factors that drive GxE.

As a second use case, we applied StructLMM to test for cell-context interactions in a large blood eQTL study. The same modelling principles enabled the identification of context-specific eQTL. Several of these interaction eQTL colocalised with GWAS variants and the marker genes of the cellular environment that underlie these GxE effects could be connected to plausible biological processes. This analysis also confirms that StructLMM can be robustly applied to analyse interaction effects driven by large numbers of environments.

Although we found that StructLMM is a robust alternative to conventional linear interaction tests, the model is not free of limitations. First, while the computational complexity of the model scales linearly with the number of individuals, thereby enabling genome-wide analyses of large cohorts, its application remains computationally more demanding than conventional LMMs. A second area for future developments is the selection of variants for GxE tests. To mitigate the cost of multiple testing, we have considered variants that have been associated with the phenotype in other studies. However, the fact that our association tests identifies novel loci if applied genome-wide suggests that this filter is not optimal. Finally, while StructLMM can in principle be used in conjunction with any environmental covariance, we have here limited the application to linear covariances. The model could be extended to account for non-linear interactions, for example using polynomial covariance functions. Such developments are a future area of work, in particular as increasingly large cohorts allow for detecting such higher order interaction effects.

**Availability of code and data.** StructLMM is available from https://github.com/limix/struct-lmm and is supported within the LIMIX framework at https://github.com/limix/limix. For tutorials and illustrations on how to use the model, see http://struct-lmm.readthedocs.io. The BIOS RNA data can be obtained from the European Genome-phenome Archive (EGA; accession/EGAS00001001077). Genotype data are available from the respective biobanks.

## Acknowledgements

The others would like to thank Christoph Lippert and Leopold Parts for helpful discussions. This research has been conducted using the UK Biobank Resource (Application Number 14069). R.M. was supported by a PhD fellowship from the Mathematical Genomics and Medicine programme, funded by the Wellcome Trust. F.P.C. & O.S. received support from core funding of the European Molecular Biology Laboratory and the European Union’s Horizon2020 research and innovation programme under grant agreement N635290. IB acknowledges funding from Wellcome (WT098051 and WT206194). The Biobank-Based Integrative Omics Studies (BIOS) Consortium is funded by BBMRI-NL, a research infrastructure financed by the Dutch government (NWO 184.021.007).

## Author contributions

R.M., F.P.C., I. B., & O.S. conceived the method. R.M., F.P.C. implemented the methods. R.M., F.P.C., M.J.B. analyzed the data. D.H. provided analyses tools. R.M., F.P.C., I.B., & O.S. interpreted results and wrote the paper.

## Online Methods

The full method section is attached as supplementary file (Supplementary methods).

## Supplementary items

### Supplementary Methods

Methods, including full derivation of the StructLMM model and analysis details.

### Supplementary figures and tables

Supplementary items.

### BIOS consortium banner

Bastiaan T. Heijmans, Peter A.C.’t Hoen, Joyce van Meurs, Aaron Isaacs, Rick Jansen, Lude Franke, Dorret I. Boomsma, René Pool, Jenny van Dongen, Jouke J. Hottenga, Marleen M.J. van Greevenbroek, Coen D.A. Stehouwer, Carla J.H. van der Kallen, Casper G. Schalkwijk, Cisca Wijmenga, Alexandra Zhernakova, Ettje F. Tigchelaar, P. Eline Slagboom, Marian Beekman, Joris Deelen, Diana van Heemst, Jan H. Veldink, Leonard H. van den Berg, Cornelia M. van Duijn, Bert A. Hofman, André G. Uitterlinden, P. Mila Jhamai, Michael Verbiest, H. Eka D. Suchiman, Marijn Verkerk, Ruud van der Breggen, Jeroen van Rooij, Nico Lakenberg, Hailiang Mei, Maarten van Iterson, Michiel van Galen, Jan Bot, Peter van’t Hof, Patrick Deelen, Irene Nooren, Matthijs Moed, Martijn Vermaat, Dasha V. Zhernakova, René Luijk, Marc Jan Bonder, Freerk van Dijk, Wibowo Arindrarto, Szymon M. Kielbasa, Morris A. Swertz, Erik W. van Zwet

